# Strategies of invertebrate osmoregulation: an evolutionary blueprint for transmuting into fresh water from the sea

**DOI:** 10.1101/2022.01.16.476502

**Authors:** John Campbell McNamara, Carolina Arruda Freire

**Author notes:** Corresponding Author, John Campbell McNamara, Departamento de Biologia, FFCLRP, Universidade de São Paulo, Ribeirão Preto 14040-901, SP, Brazil.

## Abstract

Early marine invertebrates like the Branchiopoda began their sojourn into dilute media some 500 million years ago in the Middle Cambrian. Others like the Mollusca, Annelida and many crustacean taxa have followed, accompanying major marine transgressions and regressions, shifting landmasses, orogenies and glaciations. In adapting to these events and new habitats such invertebrates acquired novel physiological abilities that attenuate the ion loss and water gain that constitute severe challenges to life in dilute media. Among these taxon-specific adaptations, selected from the subcellular to organismal levels of organization, are reduced body permeability and surface (S) to volume (V) ratios, lowered osmotic gradients, increased surface areas of interface epithelia, relocation of membrane proteins in ion-transporting cells and augmented transport enzyme abundance, activity and affinity. We examine adaptations in taxa that have penetrated into fresh water, revealing diversified modifications, a consequence of distinct body plans, morpho-physiological resources and occupation routes. Contingent on life history and reproductive strategy, numerous patterns of osmotic regulation have emerged, including intracellular isosmotic regulation in weak hyper-regulators and well-developed anisosmotic extracellular regulation in strong hyper-regulators, likely reflecting inertial adaptations to early life in an estuarine environment. Our analyses show that across sixty-four freshwater invertebrate species from six phyla/classes, hemolymph osmolalities decrease logarithmically with increasing S: V ratios. The arthropods have the highest osmolalities, from 300 to 650 mOsmoles/kg H_2_O in the Decapoda with 220 to 320 mOsmoles/kg H_2_O in the Insecta; osmolalities in the Annelida range from 150 to 200 mOsmoles/kg H_2_O, the Mollusca showing the lowest osmolalities at 40 to 120 mOsmoles/kg H_2_O. Overall, osmolalities reach a cut-off at ≈200 mOsmoles/kg H_2_O, independently of increasing S: V ratio. The ability of species with small S: V ratios to maintain large osmotic gradients is mirrored in their putatively higher Na^+^/K^+^-ATPase activities that drive ion uptake processes. Selection pressures on these morpho-physiological characteristics have led to differential osmoregulatory abilities, rendering possible the conquest of fresh water while retaining some tolerance of the ancestral medium.

## Introduction

Invertebrate organisms such as crustaceans, mollusks and annelids likely began their conquest of the freshwater biotope from their ancestral marine settings in the early Carboniferous (≈360 million years ago) (Calder, 1998; Iglikowska, 2014) or even earlier in the case of certain basal crustacean taxa like Branchiopoda (512-478 million years ago, Middle Cambrian to Early Ordovician, Sun et al., 2016) and Ostracoda (400-370 million years ago, Devonian, Martens et al., 2008; Ayhong and Huang, 2021). Some paleontological records suggest that this initial massive colonization was the result of numerous marine transgressions and regressions (Whatley, 1990; Tibert and Scott, 1999; Williams et al., 2006; Bennett, 2008; Bennett et al., 2012). Transgressional events could provide aquatic continuity, enabling the ingression of coastal and shallow water species and the eventual evolution of tolerance to environments of low or fluctuating salinity, while regressions would trap intrusive marine species that would eventually become adapted to their ensuing low salinity or freshwater setting. Whatever the process, such transmuted invaders would have been subject to selection pressures favoring the evolution of many biochemical, morphological, physiological and behavioral adaptations, including life history strategies, some probably still dependent on saline or brackish waters, that would enable eventual success in their conquest of fresh water, at least as adult semaphoronts (Havstad et al., 2015; Sharma et al., 2017; Luque et al., 2021).

At the other end of the spectrum of organization, from a cellular point of view, the optimal characteristics necessary to sustain the vibrant spark of life must be maintained to allow its inherent subcellular biochemical reactions. Early transmuters into fresh water would likely possess little ability to regulate their intracellular, and particularly extracellular fluids, given their ancestry in a marine *milieu*. In this particular environment, major ion and water concentrations are fairly stable and such organisms, like their extant relatives today, would tend to hold their body fluids in isosmoticity with their surroundings, expending little energy on anisosmotic extracellular regulation (Kirschner, 1991). Nevertheless, they would regulate the gradients and concentrations of the major ions in their intracellular fluids using energy-dependent ion transport proteins, a consequence of previous unicellular and multicellular evolution that demanded isosmotic intracellular regulation, itself a consequence of the advent, organization and composition of the cell membrane (Florkin, 1962; Kirschner, 1991; Freire et al., 2008b; Charmantier et al., 2009; Evans, 2009; Foster et al., 2010; Cuenca et al., 2021). The ability of an organism to maintain osmotic and ionic gradients against an external medium would only arise with the organization and selection of transport proteins and enzyme isoforms capable of altering the biochemical composition of these intra- and extracellular seas, or their relocation to specific areas of cell membranes or cellular compartments. Thus, only by firstly using and rearranging the existing building blocks of ion transport proteins like the sodium-potassium and the proton adenosine triphosphatases and ion anti-porters and symporters like the sodium-proton exchanger and the sodium-potassium-two chloride carrier and ion channels, would early freshwater invertebrates be able to begin to exploit their novel environment. From these underlying energy-dependent, intra- and extracellular osmotic and ionic equilibria, would spring the subsequent evolution of successfully adapted freshwater inhabitants (Florkin, 1962; Péqueux, 1995; Freire et al. 2008a,b; Evans, 2009; Anger, 2013; Freire et al., 2013; Lee et al., 2012).

While these events envision major biochemical rearrangements that may have accompanied invertebrate organisms into fresh water, much evidence suggests that many taxa have invaded this environment independently at different times, likely unrelated to marine trans- and regressions, including derived crustacean groups like freshwater Caridea and Brachyura (Pennak, 1985; Graf and Foighil, 2000; Anger, 2013). Contemplated on more restricted temporal and spatial scales, such possible routes into continental lotic and lentic biotopes could include the putative sequential occupation of habitats ranging from fully marine, intertidal waters to variable salinity estuarine habitats to stable brackish water habitats to fresh water as has been suggested for some palaemonid shrimps (Augusto et al., 2009), aquatic Brachyura (Luque et al., 2021) and other semi-terrestrial crab taxa (Morris and Van Aardt, 1998). Other possibilities include the origins of organisms in marine habitats through the intertidal zone to semi-terrestrial habitats and then into fresh water (Schubart et al., 1998; Schubart and Diesel, 1999), or from terrestrial habitats as seen in Insecta (Roer et al., 2015; Tamone and Harrison, 2015; Kefford et al., 2016).

Clearly extant freshwater inhabitants are a heterogeneous lot, exhibiting distinct sizes, shapes, body plans, structural complexity, life histories and modes of reproduction. This diversity extends to their ability to translocate between different osmotic environments, reflecting their exploitation of temporally and spatially distinct habitats. Some are diadromous, migrating between fresh and estuarine or marine waters during different life cycle stages while others are restricted entirely to fresh water, i. e., are hololimnetic (Freire et al., 2003; Evans, 2009; Anger, 2013; Griffith, 2017). Reflecting their temporally distinct transmutations from marine ancestors, many taxa like crustaceans are still dependent on the ancestral habitat and require brackish or marine waters to complete their life cycles (Augusto et al., 2007a,b, 2009; Charmantier and Anger, 2011; Anger, 2013). Further, the different routes into fresh water taken by their ancestors were not concomitant in evolutionary time and space and are likely separated by many hundreds of millions of years, and tens of thousands of kilometers (Anger, 2013; Ayhong and Huang, 2020; Luque et al., 2021). Such disparate origins have resulted in a plethora of adaptations at many levels of organization, leading to very different regulatory capabilities as illustrated by freshwater crabs and shrimps for example (Mantovani and McNamara, 2021). Our following considerations attempt to identify patterns, trends and tendencies in osmotic and ionic regulatory mechanisms in some of the main invertebrate taxa that have become established in fresh water.

### Water and the aquatic environment

The aqueous media of aquatic environments differ widely in their physico-chemical properties and in composition. While salt content is perhaps the most important characteristic from a physiological point of view, many other parameters, including ecological attributes, define ‘water’ and affect the organisms that inhabit this remarkable liquid. These parameters include its boiling and freezing points, density, surface tension and capillarity, heats of vaporization and fusion, vapor pressure, viscosity and cohesion, conductivity, dipole moments, amphoteric nature and redox reactability, solubility, and state transformations and associated energy flux. These important characteristics have led some to consider water to be a universal solvent and the solvent of life.

Natural fresh water, be it lotic or lentic, contains various dissolved minerals, ions, gases and organic compounds owing to its interactions with the atmosphere and with the substrates over which it flows or by which it is contained. These chemical entities define water’s acidity, alkalinity, permanent or temporary hardness and softness, among other properties. Considered strictly from the viewpoint of its major dissolved ions, most of the water on the earth’s surface consists of seawater, i. e., 97.4% of the global water volume of 1.38 × 10^9^ km^3^, containing (in mmol L^−1^) the cations Na^+^ 469.0, K^+^ 10.2, Mg^2+^ 52.8 and Ca^2+^ 10.3, and anions Cl^−^ 546.0 and SO_4_^2−^ 28.2. In general, the salinity of standard seawater refers to the mass of dissolved ions per kilogram of solution (i. e., ≈34.5 g kg^−1^, or parts per thousand [‰] or Practical Salinity Units). This salinity corresponds to an osmolality of around 1,050 mOsmoles/kg H_2_O. In contrast, continental fresh waters such as rivers and lakes correspond to less than 2.8% of the global water volume and contain but a thousandth or less of the ionic concentrations of seawater (in μmol L^−1^), e. g., Na^+^ 120, K^+^ 67, Mg^2+^ 50 and Ca^2+^ 86, and Cl^−^ 83, together with HCO_3_^−^, CO_3_^2-^, NO_3_^−^ and silica, and dissolved organic compounds like tannic and humic acids. In general, the salinity of fresh waters ranges from 0.2 to 0.5 g kg^−1^ and their osmolality is usually less than 5 mOsmoles/kg H_2_O.

Given this notable disparity in ionic content, and consequently, osmolality between fresh and marine waters, and the isosmotic nature of ancestral invertebrates in seawater, eventual success in becoming established in and fully occupying the limnic environment has come about with the emergence of adaptations related mainly to diminished water and ion permeabilities, a reduction in ionic and osmotic gradients, mechanisms of active ion uptake and the excretion of a severe osmotic water load. All these adaptations are found to some degree in freshwater invertebrates, whether they be of the perhaps more basal ‘leak and replace’ or the derived ‘reduce and conserve’ strategies of establishing ionic and osmotic equilibria (Kirschner, 1991; Evans, 2009). Nevertheless, such means of constituting systemic conditions for vital physiological processes in freshwater animals depend on the evolutionary appearance of transport and other proteins in specialized, interface epithelia, together with attendant structural modifications that underlie gas exchange, metabolism and energy supply, ultimately in the form of adenosine triphosphate.

### Osmotic gradients in freshwater invertebrates

The distinct degrees of osmoregulatory capability seen in extant organisms that inhabit fresh water may reflect how different invertebrate lineages have become adapted to the principal challenges of this biotope, i. e., limited salt availability and abundance of osmotically gainable water. Other abiotic parameters such as temperature, pH, alkalinity, hardness and dissolved oxygen, have not demanded such exclusive and specific responses from freshwater inhabitants. For example, adaptations to hypoxia are found in abyssal animals while adaptations to latitudinal temperature patterns are seen in aquatic continental and marine species (Willmer et al., 2005; Somero et al. 2017).

If an organism’s internal fluids are to be more concentrated than that of the surrounding medium, then attaining steady-state equilibria necessarily depends on reduced osmotic and ionic permeability of the body surfaces and, particularly, of the multifunctional gills. The exoskeleton provides this function in arthropods, which are further modified by different degrees of carcinization, i. e., thickening of the cuticle and reduction of the abdomen and pleopods, which are folded under the cephalothorax in Brachyura and Anomura. Consequently, cuticle osmotic and sodium permeabilities are respectively 36-fold and 17-fold lower in hyperosmoregulating freshwater decapods compared to marine osmoconformers or to intertidal and estuarine species that take up salt from brackish waters (Kirschner, 1991). Mean cuticle density increases progressively by 6-fold up to 2 weeks after ecdysis in the freshwater crayfish *Astacus astacus*, while apparent water permeability decreases concomitantly by ≈60% (Rasmussen and Andersen, 1996).

A survey of hemolymph osmolality in six phyla/classes, including a wide variety of 64 freshwater invertebrate species from continental streams, rivers, lakes and ponds (Figs. 1A, 1B) reveals that mean osmotic concentrations (mOsmoles/kg H_2_O) range from 60 in the Rotifera and Cnidaria (Benos and Prusch, 1972; Epp and Winston, 1977), 43 in the Bivalvia to 100 in Gastropoda (Byrne et al., 1989; Byrne and Dietz, 1997; Jordan and Deaton, 1999; Brix et al., 2011), 160-180 in Annelida (Holmstrup et al., 1999), 140 to 290 in Copepoda and Branchiopoda (Bayly, 1969; Brand and Bayly, 1971; Aladin and Potts, 1995; Lee et al., 2012), 230-330 in Insecta (e.g., Edwards, 1982; Gainey Jr, 1984; Pallarés et al., 2015; Dowse et al., 2017; Kengne et al., 2019), 295 in Amphipoda (Funck et al., 2013; Vellinger et al., 2013) to 450 in the Decapoda (Denne, 1968; Harris and Micallef, 1971; Castille and Lawrence, 1981; Moreira et al., 1983; 1988; Warburg and Goldenberg, 1984; Truchot, 1992; Freire et al., 2003; Khodabandeh et al., 2005; Gonzalez-Ortegon et al., 2016) (see all data and references in Supplementary Table S1).

**Figure 1.**
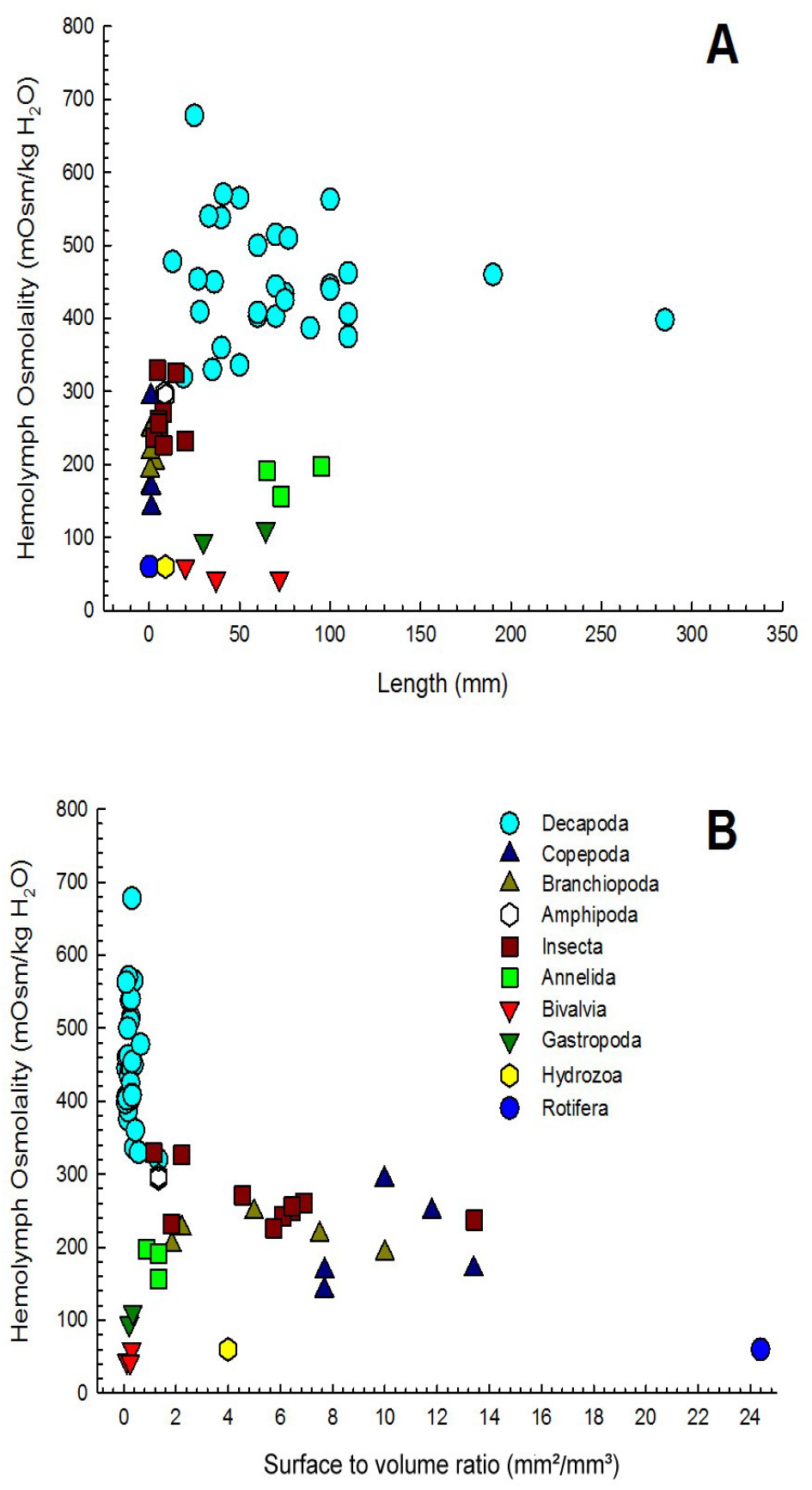
Relationships between the osmolality of the hemolymph (mOsmoles/kg H_2_O) maintained against fresh water (<0.5 ‰S, <10 mOsmoles/kg H_2_O) and (**A**) body length (mm) or (**B**) body surface to volume ratio (mm^2^/mm^3^) in 64 species of aquatic invertebrates from six phyla/classes (Crustacea, Insecta, Mollusca, Annelida, Cnidaria, Rotifera) that have successfully occupied fresh water. Raw data and references are provided in Supplementary Table S1.

Most findings concern branchiopods, copepods and decapods, given their radiation into and evolutionary success in fresh water, but also include insects and their larvae. The highest osmotic gradients against fresh water are generated by the arthropods, around 60: 1. Freshwater decapods are among the largest crustaceans and exhibit the highest hemolymph osmolalities of all invertebrates (see Supplementary Table S1 for all values), ranging from 350 to 450 mOsmoles/kg H_2_O. While they include a wide range of body lengths from 10 to 300 mm (Fig. 1A), their estimated body surface: volume (S: V) ratios are fairly constant and small, usually <0.2 mm^2^/mm^3^ (Fig. 1B). In contrast, insect larvae are much smaller, from 5 to 20 mm in body length (Fig. 1A). However, while their hemolymph osmolalities show less variation than decapods (≈100 mOsmoles/kg H_2_O), varying from 220 to 320 mOsmoles/kg H_2_O, their S: V ratios range from about 2 to 14 mm^2^/mm^3^ (Fig. 1B and Supplementary Table S1). The copepods, even smaller yet, show a similar relationship (Figs. 1A, 1B). Freshwater annelids also possess a cuticle and sustain high osmotic gradients of 36: 1 (Holmstrup et al., 1999). Considering all 64 species from six phyla/classes, hemolymph osmolality correlates negatively with S: V ratio (Pearson’s R= −0.473, P <0.001) and diminishes logarithmically with increasing S: V ratios (R^2^= 0.535). Similarly, when considering the 53 freshwater arthropods alone, hemolymph osmolality declines rapidly with increasing S: V ratios (Pearson’s R= −0.716, P <0.001) showing an inverse logarithmic relationship (R^2^= 0.800, Fig. 2 and supplementary Table S2). Thus, larger arthropods with smaller body S: V ratios can maintain greater osmotic gradients against fresh water, possibly underpinned by their higher Na^+^/K^+^-ATPase activities (see Fig. 5) and very low cuticular permeabilities.

**Figure 2.**
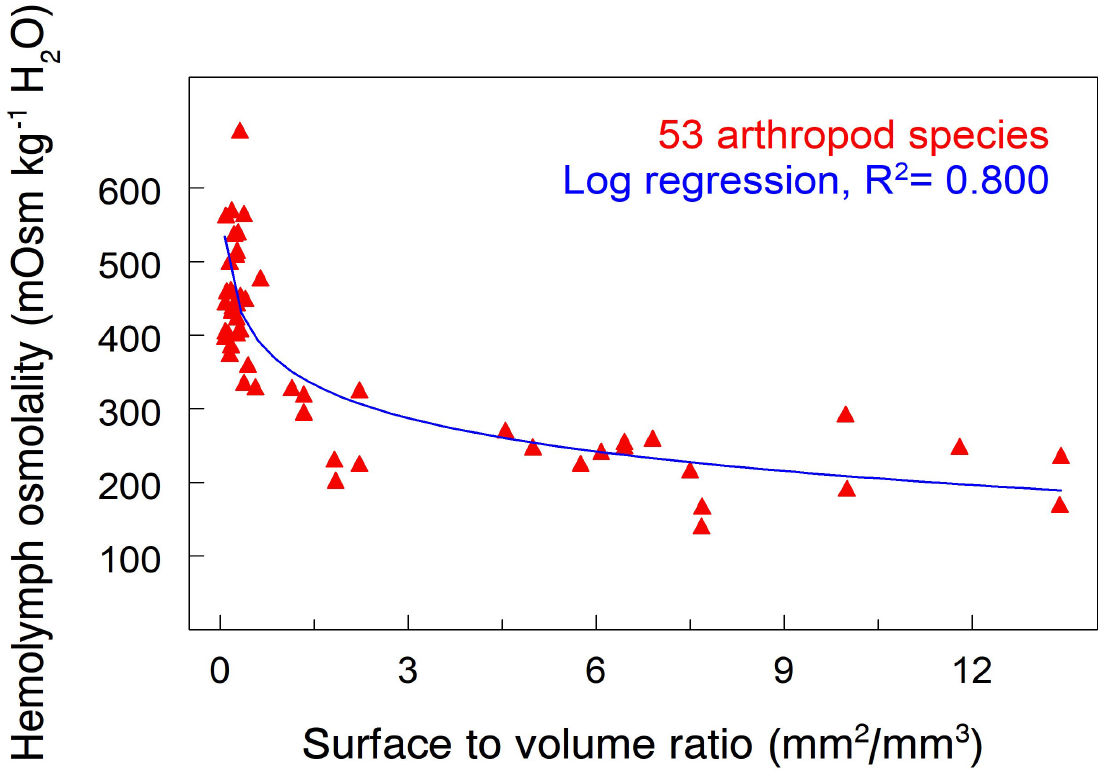
Relationship between hemolymph osmolality (mOsmoles/kg H_2_O) and surface: volume ratio (mm^2^/mm^3^) in 53 species of arthropods. Osmolality declines logarithmically with increasing surface: volume ratio (R^2^= 0.80, Pearson’s R= − 0.72, P <0.001). Raw data and references are provided in Supplementary Table S2.

Burgeoning questions arise concerning the presumptive advantages to maintaining an elevated osmotic gradient and in retaining the ability to secrete salt in some successful freshwater inhabitants like the Crustacea. Given their marine origin and the subsequent evolution of enzyme systems in cells isosmotic with seawater, cytosolic conditions ideal for sustaining the reactions of life likely would have been established early on. However, since many proteins and enzymes denature with changes in ionic concentration and pH, selection pressure might favor conserving those cellular conditions in which such essential metabolic pathways had evolved (Somero et al., 2017). Thus, during their lengthy transmutation into fresh water, once ancestral marine crustaceans have decreased their hemolymph osmolality by roughly half compared to extant marine species while holding their osmotic gradient against freshwater at around 400 mOsmoles/kg H_2_O (Faria et al., 2011). This evolutionary trade-off has been driven by mechanisms of anisosmotic extracellular and isosmotic intracellular osmoregulation that have partially conserved the primitive cytosolic environment while attenuating osmotic water gain and diffusive salt loss.

As a corollary, however, the benefit of the arthropod exoskeleton as a barrier to ion and water movements is temporally interrupted at regular intervals, since arthropods must molt to grow (Cheong et al., 2015). During pre-molt, increased osmotic permeability and resultant water influx leads to swelling and ecdysis, enabling the discontinuous growth of an ever-larger individual during the early post-molt (Freire et al., 2013). Nevertheless, molting in arthropods like freshwater crustaceans and insect larvae, requires dealing with a large, transient reduction in hemolymph osmolality and the ability to regulate cell volume (Freire et al., 2013).

Besides the Crustacea, other major freshwater taxa have been successful in confronting the challenge of inhabiting fresh water and dilute media. While such organisms cannot rely on low osmotic and ionic permeabilities owing to their diminutive size, body plans and the constraints of soft tissue structure, they have acquired the ability to produce a dynamic, steady-state equilibrium, maintaining low osmotic gradients as seen in bivalves, cnidarians and rotifers that exhibit extracellular osmolalities of from 35 to 60 mOsmoles/kg H_2_O (Fig 1A, B). Within the Mollusca, the Gastropoda appear to maintain larger osmotic gradients than the Bivalvia (Deaton, 2009), perhaps owing to their body plan in which less permeable soft tissue is exposed. The hemolymph osmolality of gastropods can exceed 100 mOsmoles/kg H_2_O (Brix et al., 2011).

### Models of invertebrate salt uptake and the Na^+^/K^+^-ATPase

While developed for the Crustacea, current models of ion uptake across the gill epithelia of freshwater shrimps and crabs likely apply to other distantly related taxa since they derive from ancient transport mechanisms in the Metazoa (McNamara and Faria, 2012; see Griffith, 2017 for models including insects and mollusks).

In two such models, ion transporters are distributed asymmetrically across the highly amplified apical and basal membranes of specialized, ion transporting, epithelial ionocytes. In weakly hyper-osmoregulating, diadromous and/or estuarine species with ion-leaky, low-resistance epithelia, the apical membrane contains a suite of antiporters that employ metabolic end products as counter ions for Na^+^ and Cl^−^, such as the Na^+^/H^+^(NH_4_^+^) and Cl^−^/HCO_3_^−^ exchangers, both driven by the metabolic hydration of CO_2_ by carbonic anhydrase (Fig. 3). The Na^+^-K^+^-2Cl^−^ symporter may be present. Such species use a ‘leak and replace’ or ‘uptake and lose’ strategy to maintain a dynamic osmotic equilibrium.

**Figure 3.**
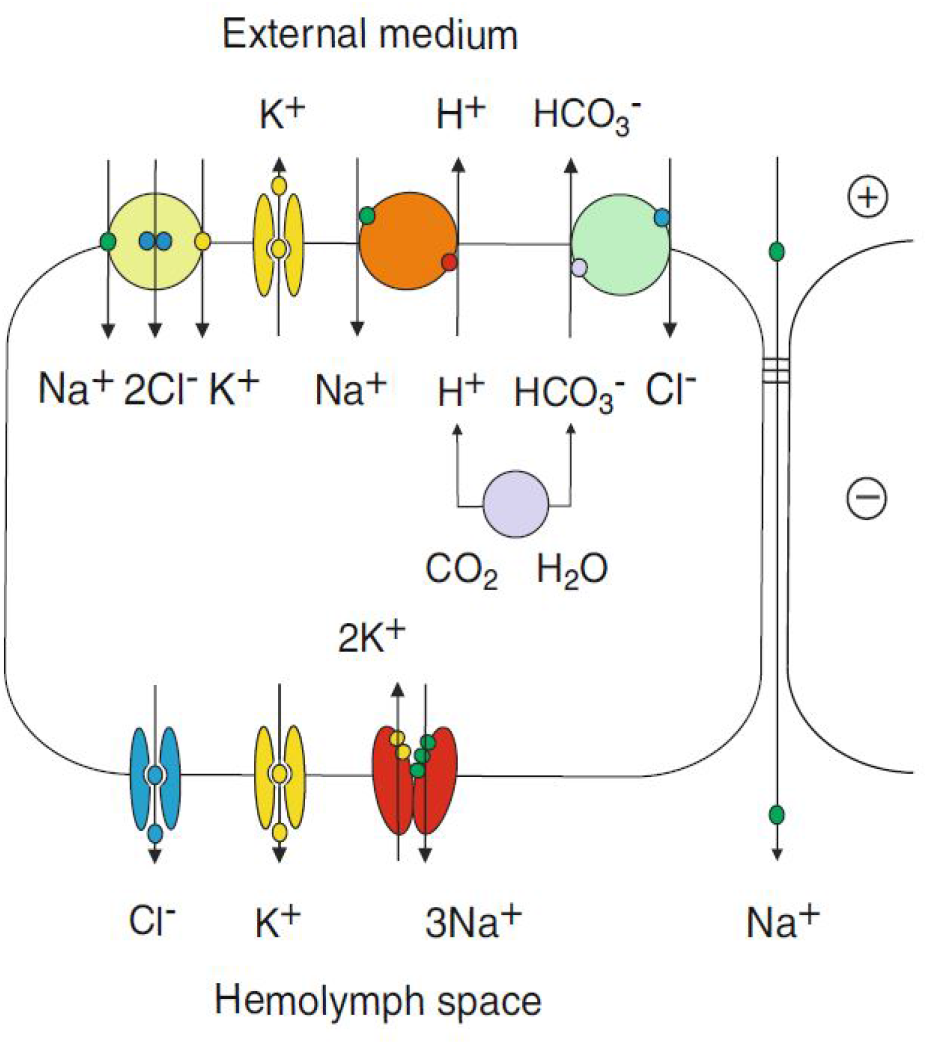
Model of coupled Na^+^ and Cl^−^ transport across a low-resistance leaky gill epithelium in weakly hyper-osmo-regulating intertidal/estuarine brachy-urans. Na^+^ and Cl^−^ flow across an apical Na^+^/K^+^/2Cl^−^ symporter driven by the inward Na^+^ gradient, supplemented by apical K^+^ channels that recycle K^+^, hyperpolarizing the apical membrane. The negative cell potential drives Cl^−^ efflux across basal Cl^−^ channels. Na^+^ also flows across an apical Na^+^/H^+^ antiporter, exiting into the hemolymph via the Na^+^/K^+^-ATPase. The outside-positive transepithelial potential drives para-cellular Na^+^ flux across the leaky epithelium. The overlying cuticle is not represented; only a single side of the gill lamella is shown (from McNamara & Faria, 2012).

In contrast, in strongly hyper-osmoregulating, hololimnetic species that exhibit tight, high-resistance epithelia, the apical Na^+^/H^+^(NH_4_^+^) antiporter has been replaced by a combination of the V(H^+^)-ATPase proton pump and Na^+^ channels through which apical Na^+^ flows down its electrical gradient into the hyperpolarized cytosol consequent to proton extrusion (McNamara and Faria, 2012, Fig. 4). This adaptation constitutes part of an ‘uptake and retain’ strategy based on low passive water and ion permeabilities.

**Figure 4.**
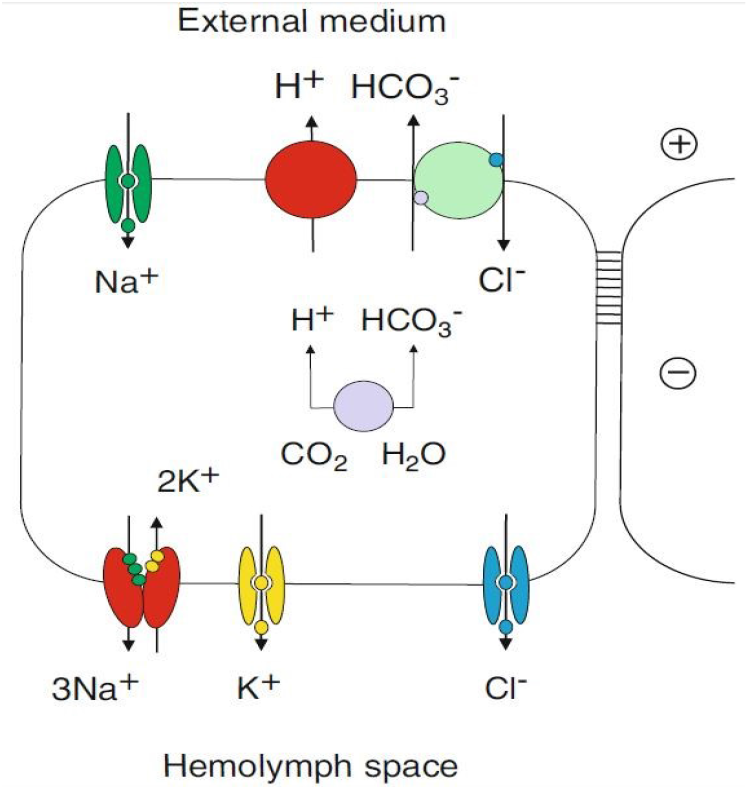
Model for uncoupled Na^+^ and Cl^−^ transport across a high-resistance gill epithelium in ‘strongly’ hyper-osmoregulating freshwater crabs. An apical V(H^+^)-ATPase pumps protons into the subcuticular space, hyperpolarizing the apical membrane, creating an electronegative apical cytosol that favors Na^+^ flux through apical Na^+^ channels down the electrical gradient. Cl^−^ is exchanged across an apical Cl^−^/HCO_3_^−^ antiporter, exiting into the hemolymph through basal Cl^−^ channels, accompanied by Na^+^ via the Na^+^/K^+^-ATPase. Paracellular ion diffusion is negligible. Only a single side of the gill lamella is shown and the overlying cuticle is not represented (from McNamara & Faria, 2012).

**Figure 5.**
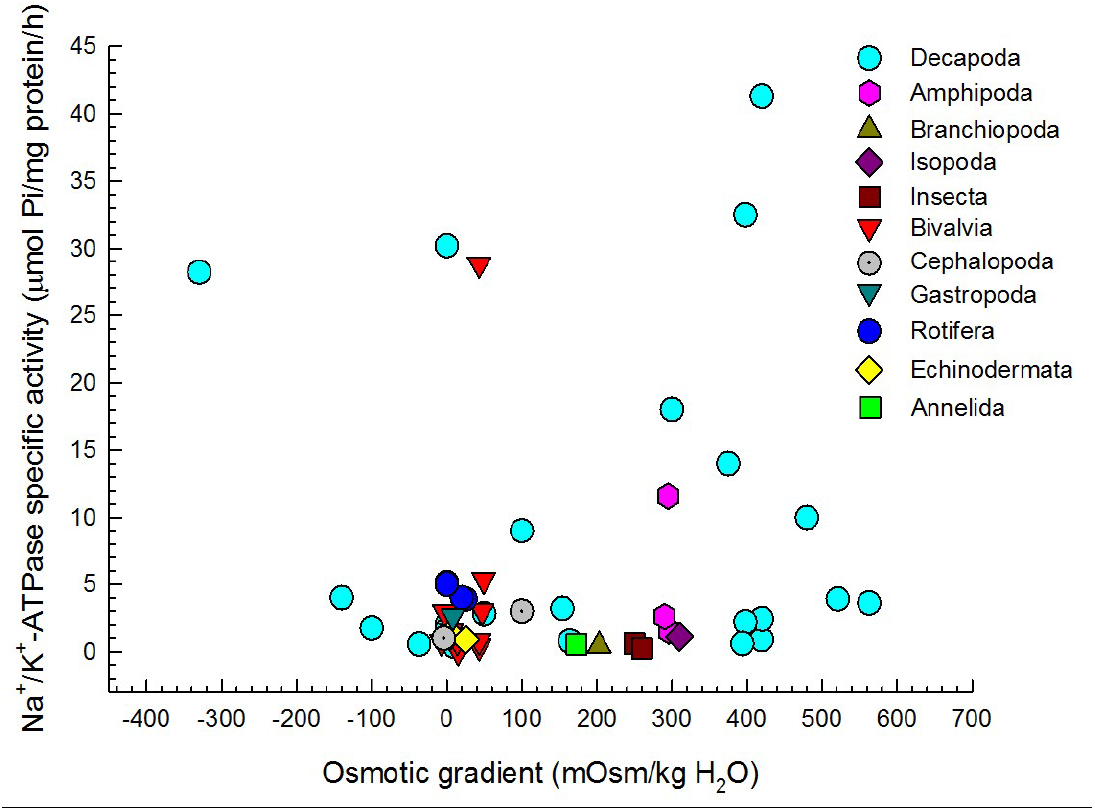
Relationship between epithelial Na^+^/K^+^-ATPase activity (μmol P_i_/mg protein/h) and osmotic gradient (mOsmoles/kg H_2_O) against the external medium in 38 species from the Crustacea, Insecta, Mollusca, Echinodermata, Annelida and Rotifera. Positive osmotic gradients reflect hyper-osmotic regulation against dilute media and are more common than negative osmotic gradients that reflect hypo-osmotic regulation against concentrated media. The absence of an osmotic gradient indicates hemolymph isosmoticity with the external medium. Raw data and references are provided in Supplementary Table S3.

In both scenarios, the basal membranes house the electrogenic Na^+^/K^+^-ATPase that counter transports 3Na^+^ into the hemolymph against 2K^+^ into the cytosol; the apically hyperpolarized cytosol drives Cl^−^ flow into the hemolymph through basal Cl^−^ channels, resulting in overall salt uptake.

Nevertheless, our analyses of 38 freshwater species from six phyla/classes dominated by the Crustacea reveal no correlation (R= 0.199, P= 0.237) between epithelial Na^+^/K^+^-ATPase activity and osmotic gradient (Δ_hemolymph/freshwater_) in hyper-osmoregulators (Fig. 5). Specific activities in the osmoregulatory tissues of many species, assayed *in vitro* from crude tissue homogenates, lie between 0.3 and 5.0 μmol P_i_ mg^−1^ protein h^−1^ while the osmotic gradient established against fresh water ranges from 40 to 560 mOsmoles/kg H_2_O. Although some crustaceans do exhibit higher activities, between 10 and 40 μmol P_i_ mg^−1^ protein h^−1^, their osmotic gradients against fresh water are unremarkable, between 300 and 400 mOsmoles/kg H_2_O. In some migratory species like *Eriocheir sinensis*, hemolymph Na^+^ gradient and gill Na^+^/K^+^-ATPase activity do correlate: the highest activities occur fresh water but decrease in brackish water where osmotic and Na^+^ gradients are reduced (Hongyu et al., 2008).

Clearly, one common evolutionary solution to maintaining the salt content of the extracellular fluid has been to diversify the suite of apical exchangers available for ion uptake across transport epithelia rather than to increase basal Na^+^/K^+^-ATPase activity, the primary transport mechanism of cytosolic Na^+^ into the hemolymph. However, using activity data from the various freshwater lineages as an indication of osmoregulatory ability requires caution. Assays *in vitro* may impart a limited view of Na^+^/K^+^-ATPase function and regulation (Moyes et al., 2021) since the enzyme plays roles other than providing the driving force for ion transport. Further, the V(H^+^)-ATPase rather than the Na^+^/K^+^-ATPase is the main driver of Na^+^ uptake from fresh water (see Fig. 4; Evans, 2009; Lee et al., 2012; Griffith, 2017).

### Extant freshwater taxa and their ancestral environment

Although most extant freshwater taxa have transmuted into dilute media from the sea, they exhibit varied relationships with their ancestral environment to which they may return at different stages of their life cycles. The higher the salinity tolerated by a freshwater species, the greater its euryhalinity, which appears to be size dependent (see Fig. 6, and Kefford et al., 2016). The Mollusca are the least euryhaline, most species surviving <10 ‰S (Hills et al., 2019). The Insecta are very diverse and include from salt-intolerant to very euryhaline species, like some chironomid Diptera that survive well in brackish water (Jonusaite et al., 2011); however, most species are modestly euryhaline (see Fig. 6, and Cañedo-Argüelles et al., 2012; Kefford et al., 2016; Le et al., 2021). The Decapoda are clearly the most euryhaline, showing an extended salinity range, some species reaching 40 ‰S, and showing some overlap with insect tolerances. However, minute rotifers and some cladocerans also can be very euryhaline (e. g., Epp and Winston, 1977; Aladin and Potts, 1995).

**Figure 6.**
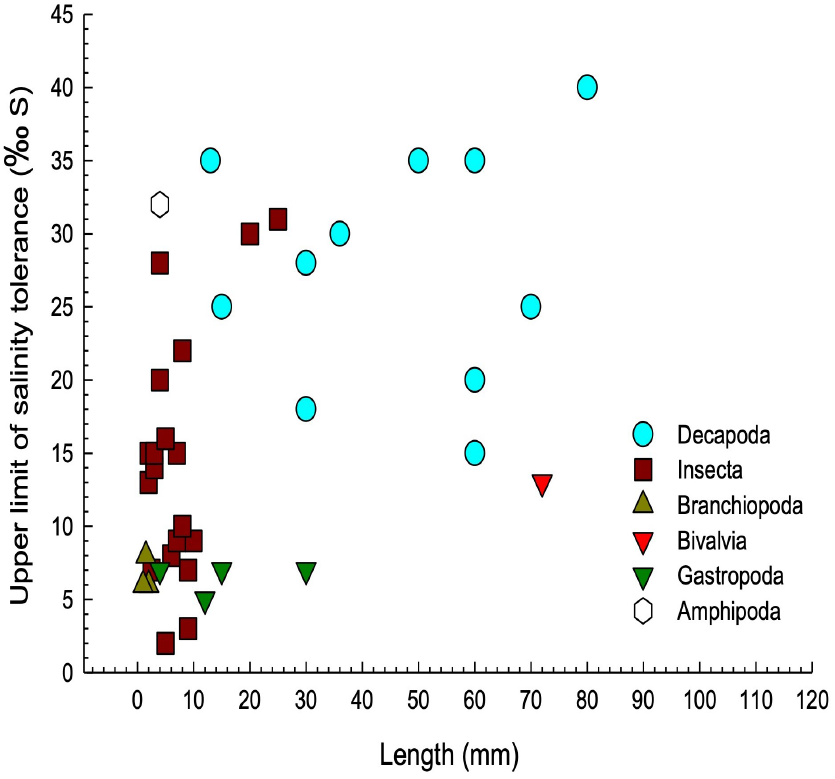
Euryhalinity as function of body length in 39 species of freshwater invertebrate taxa from the Crustacea, Insecta and Mollusca. Degree of euryhalinity is given as the approximate upper critical limit of survival at species-specific salinities compiled from various studies. Euryhalinity correlates weakly with body length. Logarithmic regression R= 0.51; Pearson’s R= 0.49, P= 0.0015. Species, raw data and references are provided in Supplementary Table S4.

Evolutionary time in fresh water is a key consideration when appreciating the history of extant freshwater invertebrates (Freire et al., 2008b; Anger, 2013; Kefford et al., 2016) and is often considered to underlie the euryhalinity and osmoregulatory plasticity of palaemonid shrimps and insects for instance (Freire et al., 2003, 2008b, 2013; Kefford et al., 2016). Nevertheless, adaptive changes in physiological traits owing to selection pressures in fresh water can manifest rapidly, reflecting strong phenotypic and genetic variability (Lee, 1999; Lee and Petersen, 2002; Lee et al., 2012). Repeated freshwater invasions have been recorded over the last 100 to 200 years in the copepod *Eurytemora affinis*, a brackish-salt marsh species complex (Lee, 1999; Lee and Petersen, 2002; Lee et al., 2012). In contrast, the trichodactylid crab *Dilocarcinus pagei* has acquired various hololimnetic adaptations over 30 to 65 million years of freshwater evolution, such as producing few large lecithotrophic eggs showing abbreviated larval development (Sternberg and Cumberlidge, 2001; Augusto et al., 2007b). Similar adaptations occur in hololimnetic palaemonid shrimps like *Macrobrachium potiuna* and *M. brasiliensis* of Mesozoic origin (60 to 100 million years ago) prior to diversification of this monophyletic genus in Neotropical fresh waters 20 million years ago (Pileggi and Mantelatto, 2010; Anger, 2013). Differently, more euryhaline diadromous species like *M. acanthurus*, *M. olfersii*, *M. rosenbergii* and *M. amazonicum* produce numerous small eggs, showing extended larval development dependent on brackish and/or coastal waters (Freire et al., 2003, 2008b; Anger, 2013). Hololimnetic anomuran aeglids show remarkable experimental salinity tolerance (McNamara and Faria, 2019), despite 25 to 75 million years’ evolution in fresh water (Pérez-Losada et al., 2004). *Aegla schmitti* tolerates 10-days salinity challenge at 25 ‰S (Pérez-Losada et al., 2004; Bozza et al., 2019; Cuenca et al., 2021) and *A. franca* at 14 to 28 ‰S (Faria et al., 2011).

Many species of diadromous freshwater Crustacea hypo-regulate their hemolymph osmolality and chloride concentrations when confronted by salinities above isosmotic (Maraschi et al. 2021). This capability enables migration between distinct halo-habitats and the exploitation of richer estuarine waters or the ancestral marine environment, suggesting an evolutionary tradeoff against the genetic and energetic costs of retaining a hyper−/hypo-regulatory apparatus. Some species still exhibit obligatory larval dependence for development on salt water that their post-larvae cannot tolerate, being obligated to migrate into fresh water, suggesting a more recent process of invasion (Augusto et al., 2007a; Charmantier and Anger, 2011). Salt secretory ability in diadromous forms like the palaemonid shrimps appears to reflect a capability advantageous in estuarine ancestors (McNamara and Faria, 2012) that has been the retained and incorporated into the migratory life cycles of more recent freshwater invaders. Why hololimnetic species have retained this ability (Moreira et al., 1983; Freire et al., 2003) is far less clear since they do not encounter salt water during their life cycle. Some of the molecular and cellular machinery necessary to effect hemolymph hyper-osmotic and ionic regulation is common to mechanisms of both salt uptake and secretion (McNamara and Faria, 2012). Ion transport proteins like the Na^+^/K^+^- and V(H^+^)-ATPases and Na^+^/H^+^(NH_4_^+^) and Cl^−^/HCO_3_^−^ antiporters are critical to ion uptake in fresh water while the Na^+^/K^+^-ATPase and the Na^+^-K^+^-2Cl^−^ symporter underlie salt secretion. The interdependence of hyper−/hypo-osmotic and ionic regulatory abilities on such proteins appears to have led to the inertial retention of an inapposite functional ability in wholly freshwater species. Alternative strategies of dealing with salt challenge have appeared in freshwater Brachyura and Anomura that simply maintain isosmoticity above their isosmotic points and rely on isosmotic intracellular regulation for volume adjustments up to their lethal salinity limits (Augusto et al., 2007; Faria et al., 2011; Mantovani and McNamara, 2021).

Thus, while overall evolutionary time in fresh water is fundamental, distinct degrees of adaptation do occur within a given taxon, reflected in different degrees of species’ euryhalinity (see variability in Fig. 6, and Supplementary Table S4). The Malacostraca include many freshwater lineages like the decapods that tolerate their ancestral medium well, likely aided by their large size (see Fig. 6). However, other taxa that also have transmuted into fresh water are not as euryhaline. Branchiopods have a much older evolutionary history, around 500 million years in fresh water (Sun et al., 2016), and separated from Malacostraca earlier than did the Insecta (Legg et al., 2013; Tamone and Harrison, 2015). The Hexapoda, while phylogenetically closer to Malacostraca (Legg et al., 2013, Tamone and Harrison, 2015) have their origin in a common Pan-crustacean ancestor but have evolved a tracheal breathing system, compatible with their terrestrial environment (Kefford et al., 2016) and only their larvae are truly aquatic. The Pan-crustacean cuticle facilitates large osmotic gradients against fresh water (Roer et al., 2015; Kefford et al., 2016) as seen in some insect larvae and the decapod Malacostraca. In contrast, the Ephemeroptera, Plecoptera and Trichoptera are the least euryhaline insect orders and are impacted by anthropogenic salinization at 3-5 ‰S, as are entire freshwater invertebrate communities (Cañedo-Argüelles et al., 2012; Kefford et al., 2016; Le et al., 2021; see Supplementary Table S4).

### Conclusions and perspectives

A final issue concerns evaluation of the physiological adaptedness of freshwater invertebrates to their environment, given their taxonomic diversity and their distinct osmoregulatory abilities. Several strategies seem to have arisen independently and may be useful indicators of adaptation or of evolutionary processes in progress. The elevated salinity tolerances of some euryhaline Malacostraca suggest a more recent transmutation into fresh water from their ancestral marine environment compared to stenohaline freshwater taxa, and thus these may still be adapting (Augusto et al., 2007, 2009; Faria et al., 2011; Maraschi et al., 2021). The retention of unexpected salinity tolerance in very old hololimnetic dwellers like the Anomura and Brachyura is remarkable; these rely on isosmoticity rather than on hypo-osmotic regulation when confronted by experimental salt challenge (Faria et al., 2011; Mantovani and McNamara, 2021) a characteristic of some marine species. The diminished hemolymph osmotic gradient against fresh water seen in Rotifera, Cnidaria, and Mollusca would diminish energy expenditure on ion uptake and seems to be adaptive. However, the elevated osmolality of most decapod Crustacea does not fit well with this scheme, particularly the hololimnetic Brachyura and Anomura, and the many diadromous Caridea that are dependent on salt water for larval development. Maintaining a low hemolymph isosmotic point compared to marine ancestors in those euryhaline taxa that are salt tolerant also suggests freshwater adaptedness (Augusto et al., 2009) as does a low inflection point with increasing hemolymph osmolality. While there are caveats and exceptions to each of these strategies, some physiological characteristics do seem to reflect the differential adaptation of many freshwater taxa to this once hostile environment.

## Acknowledgments

We gratefully acknowledge past and ongoing financial support from the Fundação de Amparo à Pesquisa do Estado de São Paulo (FAPESP #2007/04870-9, #2010/17534-0, #2011/22537-0, #2012/06620-8, #2013/23906-5, #2015/00131-3 to JCM) and the Conselho Nacional de Desenvolvimento Tecnológico e Científico (CNPq #303282-84, #304316/2003-2, #304174/2006-8, #300662/2009-2, #300564/2013-9, #303613/2017-3 and #305421/2021-2 to JCM and #405206/2016-0 and #307760/2019-7 to CAF) in the form of research grants and Excellence in Research Scholarships that have financed this line of investigation over the years.

## Data availability

The literature data supporting the findings in this article are available in the article itself and in its referenced online supplementary material.

